# Broad-Range Virus Detection and Discovery Using Microfluidic PCR Coupled with High-throughput Sequencing

**DOI:** 10.1101/2020.06.10.145052

**Authors:** Ying Tao, Clinton R. Paden, Krista Queen, Jing Zhang, Eishita Tyagi, Suxiang Tong

## Abstract

There is a need for a comprehensive and sensitive method to test for a broad range of viral pathogens in samples without any identifiable pathogen detected. Real-time PCR assays are sensitive and rapid, but their specificity limits their utility in detecting divergent agents. Shotgun high-throughput sequencing methods provide unbiased sequence identification, however, they have limited sensitivity and require complex analyses. In order to meet the need for a sensitive, high-throughput virus detection and discovery platform with good sensitivity, we combine two existing technologies, broadly-reactive consensus-degenerate pan-viral group PCR and the MiSeq sequencer (Illumina), using the Access Array (Fluidigm), a commercially-available microfluidic PCR system. Pan-viral group primers target conserved regions of virus taxonomic groups and can amplify known and potentially novel species. The Access Array employs dozens of these assays in parallel, which are then sequenced all at once on the MiSeq. In this study, we run a respiratory panel of pan-viral group PCR assays using AA-PCR-Seq. We validate the panel on a collection of representative human and animal samples, comparing it to qPCR and shotgun next-generation sequencing (NGS). AA-PCR-Seq provides a robust, straightforward method for screening large numbers of samples for virus detection and discovery.

## 1. Introduction

Laboratory diagnosis is central to responding to health threats. Molecular-based pathogen detection systems have tremendous potential for rapidly identifying and characterizing known and novel pathogens in a complex microbial community. A number of sensitive targeted and multipathogen detection platforms have been developed, but in a significant fraction of disease cases, the etiology remains unknown (e.g. 20%-50% of acute lower respiratory tract illness, >50% of encephalitis, 35%-55% of gastroenteritis) (Centers for Disease Control URDO Working Group, 2012; Granerod et al., 2010a; Granerod et al., 2010b; Lynch et al., 2006; Wikswo et al., 2015). Thus, it is critical to have a sensitive, straigtforward pathogen identification and discovery system that can detect a very wide range of pathogens, including those not yet known.

Agent-specific quantitative, or real-time, PCR (qPCR) is an indispensable technology in molecular, culture-independent virus detection (Langley et al., 2015). When designed properly, qPCR assays are sensitive, specific, and easy to interpret. However, qPCR is limited by its narrow breadth of detection and scalability. Strategies to combat these limitations have typically included multiplexing assays in the same reaction tube, or miniaturizing assays in proprietary systems, such as the Taqman Array Card (Kodani et al., 2011). These solutions come with tradeoffs in sensitivity and flexibility and they do not address the limitation that agent-specific PCR assays cannot detect novel or divergent infectious agents.

In order to detect a broad range of known and novel microorganisms, next generation sequencing has been employed in efforts to identify all DNA sequences in a sample (Pallen, 2014). Shotgun sequencing has been effective at discovering novel pathogens (Chiu, 2013; Finkbeiner et al., 2009) and diagnosing individual unknown clinical cases (Naccache et al., 2014; Pallen et al., 2010). However, due to the low sensitivity and overall high cost, the technique is not ready for routine use in diagnosis or for large studies of potential infectious disease. The sensitivity is low because the ratio of pathogen nucleic acid relative to host can be very small for many primary or clinical samples. Many millions of sequence reads must be collected to find relatively rare pathogen sequences, which raises sequencing costs and computational effort for data analysis.

Pan-viral group PCR using consensus-degenerate primers has proven very useful in identifying known and novel viral (Rose et al., 2003; Rose et al., 1998; Staheli et al., 2011; Tong et al., 2008), including many novel orthomyxoviruses, rhabdoviruses, paramyxoviruses, and adenoviruses in (Gloza-Rausch et al., 2008; Ng et al., 2013; Quan et al., 2010; Tao et al., 2013; Tong et al., 2012; Wacharapluesadee et al., 2015; Yuan et al., 2014; Zlateva et al., 2011). Consensus-degenerate primers target conserved domains within a virus family or genus and are based on amino acid similarity. This strategy increases the likelihood that the primers will amplify both known and potentially novel members of that virus family or genus (Rose, 2005; Tong et al., 2008). Because these are PCR assays, the sensitivity for pathogen discovery is quite high in comparison to shotgun sequencing by NGS.

Pan-viral group PCR amplicons must be sequenced to make an identification, which gives more detailed genetic information than signal-based qPCR assays. However, generating, cleaning up, and sequencing these amplicons on a capillary DNA sequencer is time-consuming and expensive, especially as the number of samples and assays increases. Here, we present AA-PCR-Seq, a virus detection and discovery method that combines the direct, sensitive approach of pan-viral group PCR with high-throughput approaches afforded by the Access Array microfluidic system (Fluidigm) and Illumina sequencing. AA-PCR-Seq provides a substantial advantage in sensitivity, cost, and time-to-answer for large-scale pathogen detection and discovery studies, compared to shotgun either NGS or pan-viral group PCR alone. In this report, we use a panel of respiratory virus pan-viral group PCR assays as proof of concept to demonstrate and validate the AA-PCR-Seq method. We compare the sensitivity and specificity of AA-PCR-Seq to shotgun NGS, as well as to conventional pan-viral group PCR assays and qPCR, where possible, on a variety of virus isolates, clinical samples, and primary animal samples.

## 2. Materials and Methods

### 2.1. Viral samples and sample viral RNA

Reference virus samples, clinical specimens and bat samples used in this study are described in Table 1. Clinical specimens with known infection were selected from previous disease etiology studies and from public health responses, and de-identified. The Institutional Animal Care and Use Committee (IACUC) of the Centers for Disease Control and Prevention approved all protocols related to the animal experiments in this study.

**Table 1.**
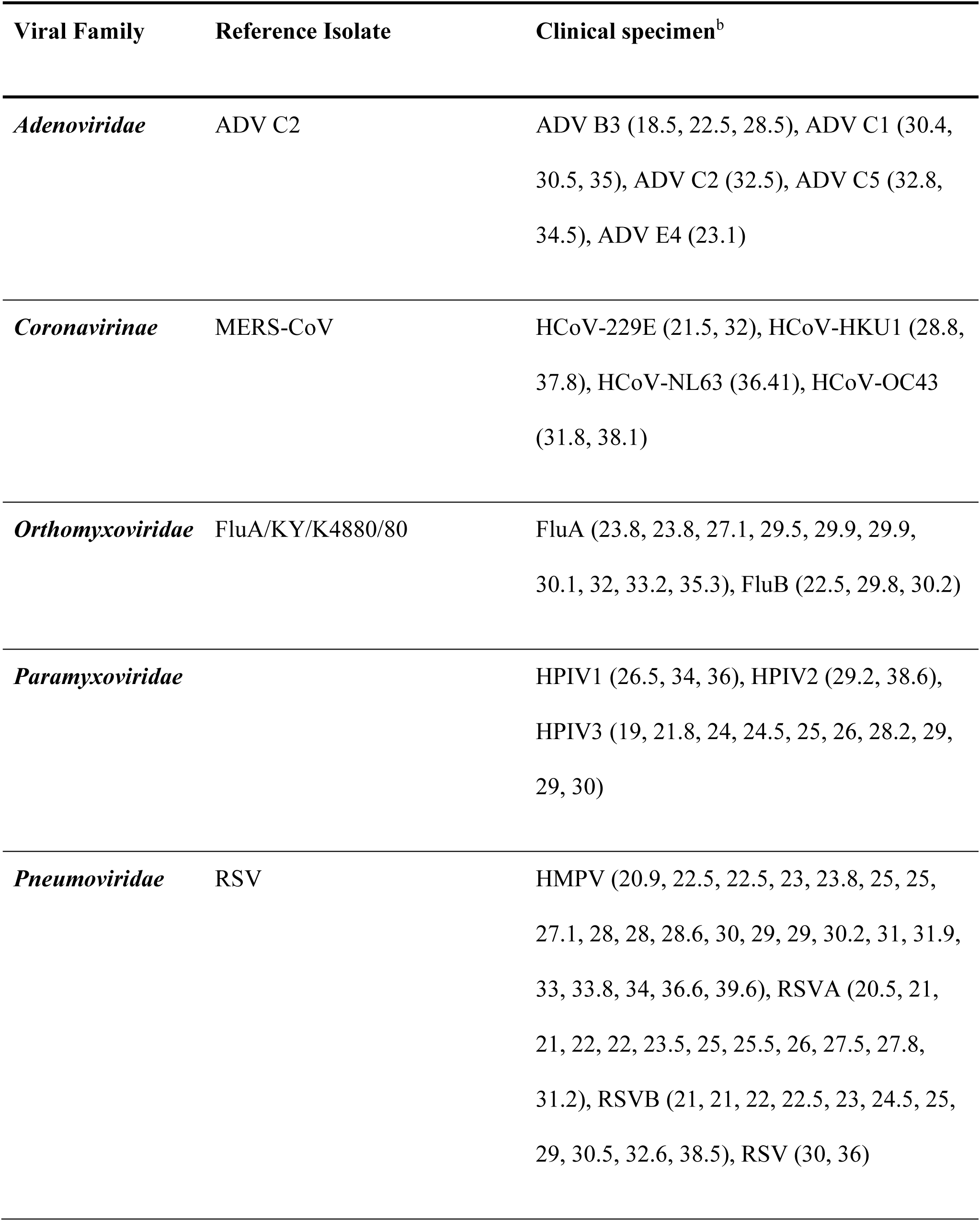
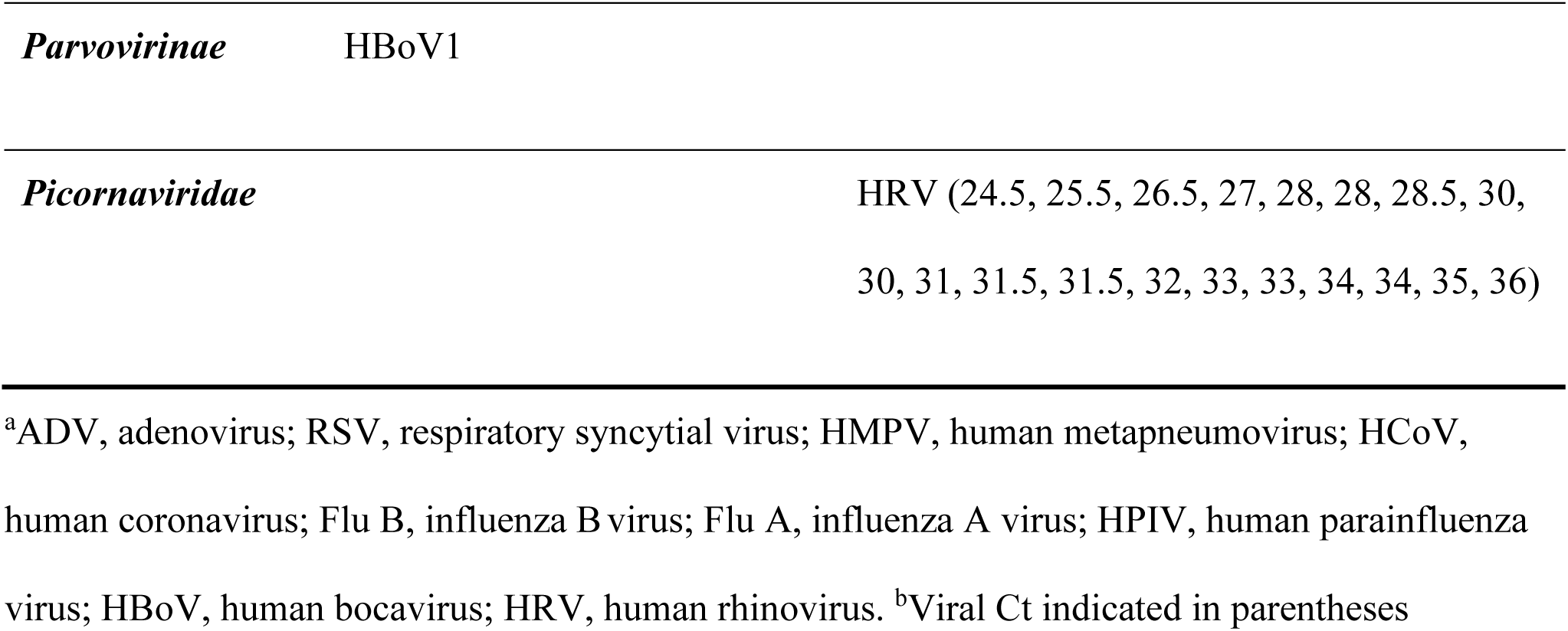
Reference and clinical specimens^a^.

RNA was extracted from 100 µL of each sample using the QIAamp viral RNA kit (Qiagen) according to the manufacturer’s instructions. The RNA was eluted from the column in 50 µL of RNase-free water and stored at −80°C.

### 2.2. Quantitative PCR and conventional pan-viral group PCR

qRT-PCRs were performed using Ambion AgPath-ID One Step RT-PCR (Thermo Fisher) according to Centers for Disease Control and Prevention-developed singleplex real-time reverse transcription PCR (rRT-PCR) assays for respiratory virus detection (Dare et al., 2007; Fry et al., 2010; Heim et al., 2003; Lu et al., 2006; Lu et al., 2008; Morgan et al., 2013; O’Flaherty et al., 2018). The pan viral-group PCRs were run using previously published protocols (Kaur et al., 2008; Schatzberg et al., 2009; Tong et al., 2008; Tong et al., 2009; Tong et al., 2010). Briefly, a nested or semi-nested set of PCRs was performed using each assay and visualized on a 2% agarose gel. PCR products were gel purified and sequenced on an ABI 3100xl capillary sequencer.

### 2.3. Random PCR-based pre-amplification

To pre-amplify samples, we used a modified “Sequence-Independent Single Primer Amplification” (SISPA) protocol (Wang et al., 2003). Nucleic acid was reverse transcribed using the SuperScript III First Strand Synthesis kit (Life Technologies) and primer A2 which contains a known sequence followed by a random nonamer (GTTTCCCAGTCACGATANNNNNNNNN). Second strand synthesis was performed using Sequenase 2.0 (ABI) and primer A2. Subsequently, the material was amplified by PCR using Primer B2 (GTTTCCCAGTCACGATA), which is homologous to the known sequence from A2. The round B product is purified using Agencourt AMPure XP beads (Beckman Coulter).

### 2.4. Pan-viral group PCR by Fluidigm Access Array and Illumina MiSeq sequencing analysis (AA-PCR-Seq)

The pan-viral group PCR for common respiratory virus group were previously described (Kaur et al., 2008; Schatzberg et al., 2009; Tong et al., 2008; Tong et al., 2009; Tong et al., 2010). For use on the Access Array, the pan-viral group PCR primers were extended at the 5’ end with linker sequence CS1 (ACACTGACGACATGGTTCTACA) on the forward primer and the linker sequence CS2 (TACGGTAGCAGAGACTTGGTCT) (Fluidigm) on the reverse primer. The CS1/CS2 sequences are used to prime a subsequent PCR which adds Illumina adapter sequences and index sequences, performtd according to the manufacturer’s instructions (Fluidigm).

Each sample well of the Access Array was loaded with 4 µL of the following reaction: 2 µL 2X OneStep buffer, 0.032 µL 50mM MgSO4, 0.12 µL DMSO, 0.2 µL 10X Loading buffer (Fluidigm), 0.04 µL RNase inhibitor (Roche), 0.16 µL SuperScriptIII RT/ Platinum Taq (Life Technologies), and 1.4 µL pre-amplified sample (2.3). Each sample/buffer mixture is loaded into three separate inlets in order to improve detection sensitivity. Each assay port was loaded with premixed PCR primers diluted to 20 µM in 1X Loading buffer (Fluidigm). Each Access Array mixes and carries out 2304 individual PCR reactions simultaneously. Reverse transcription was conducted on the Access Array at 50°C for 30 min and PCR cycling conditions of: 95°C 2 min, then 50 cycles of 15s@94°C, 30s@50°C, 30s@72°C. The 2-Primer Amplicon Tagging Protocol from the manufacturer was used (Fluidigm).

Following pan-viral group PCR on the Access Array, 10 µL of the amplicons, pooled by sample, are collected from the original sample inlets. A second PCR reaction is then performed using 1:100 diluted amplicons to add Illumina sequence-specific adaptors and sample barcodes. All barcoded samples are pooled and mixed with a 5% PhiX DNA spike in and sequenced on an Illumina MiSeq instrument using v3 PE600 reagents. Custom sequencing primers are used, per Fluidigm recommendation, in order to not sequence the linker sequences. Sequencing primers FL1 and FL2 (Fluidigm) are added to the Illumina sequencing primers, mixed, and replaced into the default primer wells on the reagent cartridge. Samples are pooled such that there are 100,000-400,000 reads obtained per sample.

### 2.5. Shotgun sequencing

Samples were first pre-amplified using SISPA (described in 2.3). Libraries were generated using the NEBNext Ultra II DNA library prep kit (NEB) and dual indexed. They were then multiplexed, spiked with 1% PhiX DNA, and run on a MiSeq with the goal of obtaining 3 million read pairs per sample. MiSeq v3 PE600 reagents were used, according to the manufacturer’s recommendation.

### 2.6. Data analysis and bioinformatics pipeline

Data was analyzed with an internally-developed pipeline (“PAIDIRT”). Briefly, residual sequencing primers are trimmed off based on homology and reads are trimmed for quality. Then, a further 26bp is removed from each end to insure the pan-viral group PCR primers are removed. Sequences are then aligned to a database of virus sequences using bowtie2 (Langmead and Salzberg, 2012) and consensus sequences are generated. BLASTN (Altschul et al., 1997) is used to correct the taxonomic assignment of each consensus sequences and obtain match quality information. The number of supporting reads, length of consensus sequence, percent match to the reference, consensus sequence, and taxonomy, including best match, are returned for each virus group. In order to call a result positive, it must be supported by 10X the number of any reads in the non-template control, in order to account for the phenomenon of index hopping. Further, it must have a match length of >80% the length of the targeted region in order to discount short alignments that may occur.

## 3. Results

### 3.1. Validation of AA-PCR-Seq using virus isolates

In order to evaluate the relative sensitivity of AA-PCR-Seq compared to shotgun sequencing, we chose of a panel of five representative virus isolates (Tables 1 and 2). The panel consists of two DNA viruses: Adenovirus C (AdVC) and Human Bocavirus (HBoV); and three RNA viruses: Respiratory Syncytial Virus (RSV), Influenza A (Flu A), and Middle East Respiratory Syndrome Coronavirus (MERS-CoV). For each virus, we made a series of 10-fold dilutions of nucleic acid in a constant background of human RNA. Each dilution was subjected to random PCR pre-amplification (described in 2.3) followed by either AA-PCR-Seq (2.4) or shotgun NGS (2.5). Forty-eight triplicate reactions were (144 total) were pooled for each MiSeq sequencing run. For shotgun NGS, we aimed to collect at least three million read pairs per sample in order to sequence 8 samples per MiSeq run with a moderate, but not excessive sequencing effort. We also performed conventional pan-viral group PCR and qPCR on each of these viruses (Table 2). To call samples positive for shotgun NGS, there must be at least a 10-fold enrichment in classified reads in the sample, compared to a no-template control and at least 10 classified reads. For AA-PCR-Seq, the reads must generate a consensus sequence that covers at least 80% of the projected size of the target amplicon and the number of classified reads must similarly be 10-fold higher than the no-template control sample.

**Table 2.** Comparison of assay sensitivity using five representative reference virus samples.

| Samples (10_fold dilutions) | qPCR results (C<sub>t</sub> value) |  | Results<sup>a</sup> (Pos or Neg) |  |  |
| --- | --- | --- | --- | --- | --- |
|  | Virus target | Host RnaseP target | Conventional<br>pan-viral<br>group PCR | AA-PCR-Seq | Shotgun<br>NGS |
| Adenovirus C (10<sup>0</sup>) | 26.2 | 29.2 | Pos | Pos | Pos |
| Adenovirus C (10<sup>1</sup>) | 29.8 | 29 | Pos | Pos | Neg |
| Adenovirus C (10<sup>2</sup>) | 33.2 | 29.4 | Pos | Pos | Neg |
| Adenovirus C (10<sup>3</sup>) | 36.9 | 29.5 | Neg | Neg | Neg |
| Adenovirus C (10<sup>4</sup>) | No Ct | 30 | Neg | ND | ND |
| Respiratory Syncytial Virus (10<sup>0</sup>) | 25.7 | 30 | Pos | Pos | Pos |
| Respiratory Syncytial Virus (10<sup>1</sup>) | 29 | 29.3 | Pos | Pos | Pos |
| Respiratory Syncytial Virus (10<sup>2</sup>) | 32.2 | 28.4 | Pos | Pos | Pos |
| Respiratory Syncytial Virus (10<sup>3</sup>) | 37.2 | 30 | Pos | Neg | Neg |
| Respiratory Syncytial Virus (10<sup>4</sup>) | No Ct | 29.5 | Neg | ND | ND |
| Influenza A (10<sup>0</sup>) | 25.5 | 24.5 | Pos | Pos | Pos |
| Influenza A (10<sup>1</sup>) | 28.1 | 26.9 | Pos | Pos | Pos |
| Influenza A (10<sup>2</sup>) | 32 | 28.4 | Neg | Pos | Neg |
| Influenza A (10<sup>3</sup>) | 35.5 | 28.6 | Neg | Pos | Neg |
| Influenza A (10<sup>4</sup>) | 39.5 | 28.8 | Neg | Neg | ND |
| Human Bocavirus 1 (10<sup>0</sup>) | 24 | 29.8 | Pos | Pos | Pos |
| Human Bocavirus 1 (10<sup>1</sup>) | 28.0 | 30.3 | Pos | Pos | Neg |
| Human Bocavirus 1 (10<sup>2</sup>) | 31.2 | 30.3 | Pos | Neg | Neg |
| Human Bocavirus 1 (10<sup>3</sup>) | 34 | 30.3 | Neg | Neg | Neg |
| Human Bocavirus 1 (10<sup>4</sup>) | 36.2 | 30 | Neg | Neg | ND |
| MERS coronavirus (10<sup>0</sup>) | 25.2 | 29.5 | Pos | Pos | Pos |
| MERS coronavirus (10<sup>1</sup>) | 28.4 | 29.5 | Pos | Pos | Pos |
| MERS coronavirus (10<sup>2</sup>) | 32 | 29.5 | Pos | Pos | Pos |
| MERS coronavirus (10<sup>3</sup>) | 35 | 29.5 | Pos | Pos | Neg |
| MERS coronavirus (10<sup>4</sup>) | 38.5 | 29.5 | Neg | Neg | ND |

<sup>a</sup> Pos, positive; Neg, negative; ND, not done.

For AdV, HBoV, FluA, and MERS-CoV, AA-PCR-Seq was capable of detecting lower titers of virus than shotgun sequencing. There was a 2-log increase in sensitivity for detecting AdV and influenza virus, while there was a 1-log increase in sensitivity for detecting HBoV and MERS-CoV by AA-PCR-Seq compared to the ones by shotgun sequencing (Table 2). RSV detection was equivalent between the methods, however there are 10-100 fold more reads detected by AA-PCR-Seq compared to shotgun sequencing, making the positive call more robust. As expected, the higher-volume conventional pan-viral group PCR was able to detect lower starting titers for some viruses tested than AA-PCR-Seq or shotgun sequencing (Table 2).

### 3.2. Detection of viruses from clinical samples

We used a set of de-identified clinical respiratory specimens to evaluate the performance of AA-PCR-Seq in a real-world setting. Each of these specimens was previously tested and found qPCR-positive for one or more for viral agents. In total, the specimens were positive for a total of 112 respiratory viruses by qPCR, with Ct values varying from 18 to 39 (Table 1).

We tested each of these clinical specimens by AA-PCR-Seq, conventional pan-viral group PCR, and shotgun NGS. Results are shown as percent agreement with qPCR testing (Figure 1A). Conventional pan-viral group PCR had 83% concordance to qPCR, AA-PCR-Seq had 79% concordance, and shotgun NGS had the lowest rate of agreement (59%) (Figure 1A). The same trend was seen for the various viruses detected, with shotgun NGS being the least sensitive (Figure 1B). Grouping the samples by Ct, we observed that the AA-PCR-Seq and conventional pan-viral group PCR concordance rate begins to decline only beyond Ct range of 32 (Figure 1C). There is 100% agreement between AA-PCR-Seq and conventional pan-viral group PCR at Ct ≤ 29 (Figure 1C). Importantly, AA-PCR-Seq was able to detect more qPCR-confirmed targets than shotgun NGS where the Ct was above 29. Above Ct 29, shotgun NGS detects 10/52 (19%), compared to 30/52 (58%) by AA-PCR-Seq.

**Figure 1.**
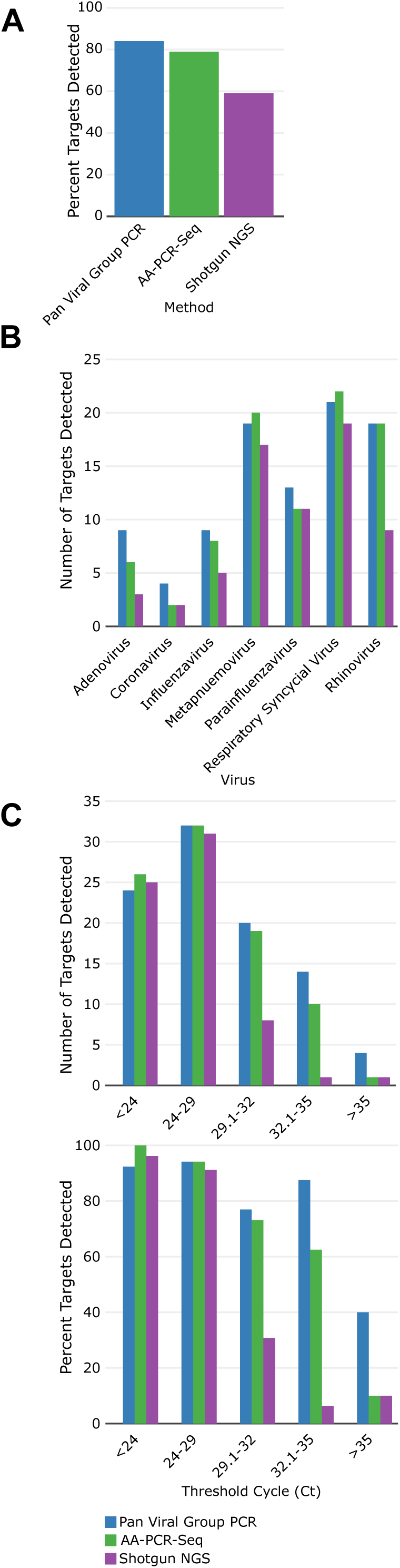
Evaluation of AA-PCR-Seq, conventional pan-viral group PCR, and shotgun NGS using clinical samples, compared to qPCR. **(A)** Overall virus detection rate, compared to qPCR, by percentage (n=112); **(B)** Rate of detection, by respiratory virus group and assay type; **(C)** Rate of detection, grouped by viral Ct value, compared to qPCR.

### 3.3. Detection of zoonotic viruses from bats

In order to test the capacity of AA-PCR-Seq to detect divergent viruses, we assembled a set of rectal and nasal swabs from various bat species which had been collected and tested as part of prior zoonotic surveillance studies. This sample set was comprised of 34 old world and new world bats from more than 21 species, from Kenya, the Democratic Republic of Congo, Democratic Republic of Georgia, and Guatemala (Table 3). Using AA-PCR-Seq, we identified 55 viruses from 4 virus families and 9 viral genera. There was an adenovirus with 95% identity based on the amplicon sequence to its nearest neighbor, 23 alphacoronaviruses with identities ranging from 80-100%, 6 betacoronaviruses with identities from 83-99%, 4 gammacoronaviruses with identities from 92-100%, 3 deltacoronaviruses with identities from 87-93%, 12 dependoparvoviruses with identities from 84-100%, one protoparvovirus with 85% identity, 4 morbiliviruses with identities from 83-92%, 1 rubulavirus with 92% identity to their nearest neighbors (Table 3). While many have divergent nucleotide sequences in the amplicon regions, additional genome sequence for each detected virus are needed to determine if they are variants, new strains, or novel species.

**Table 3.**
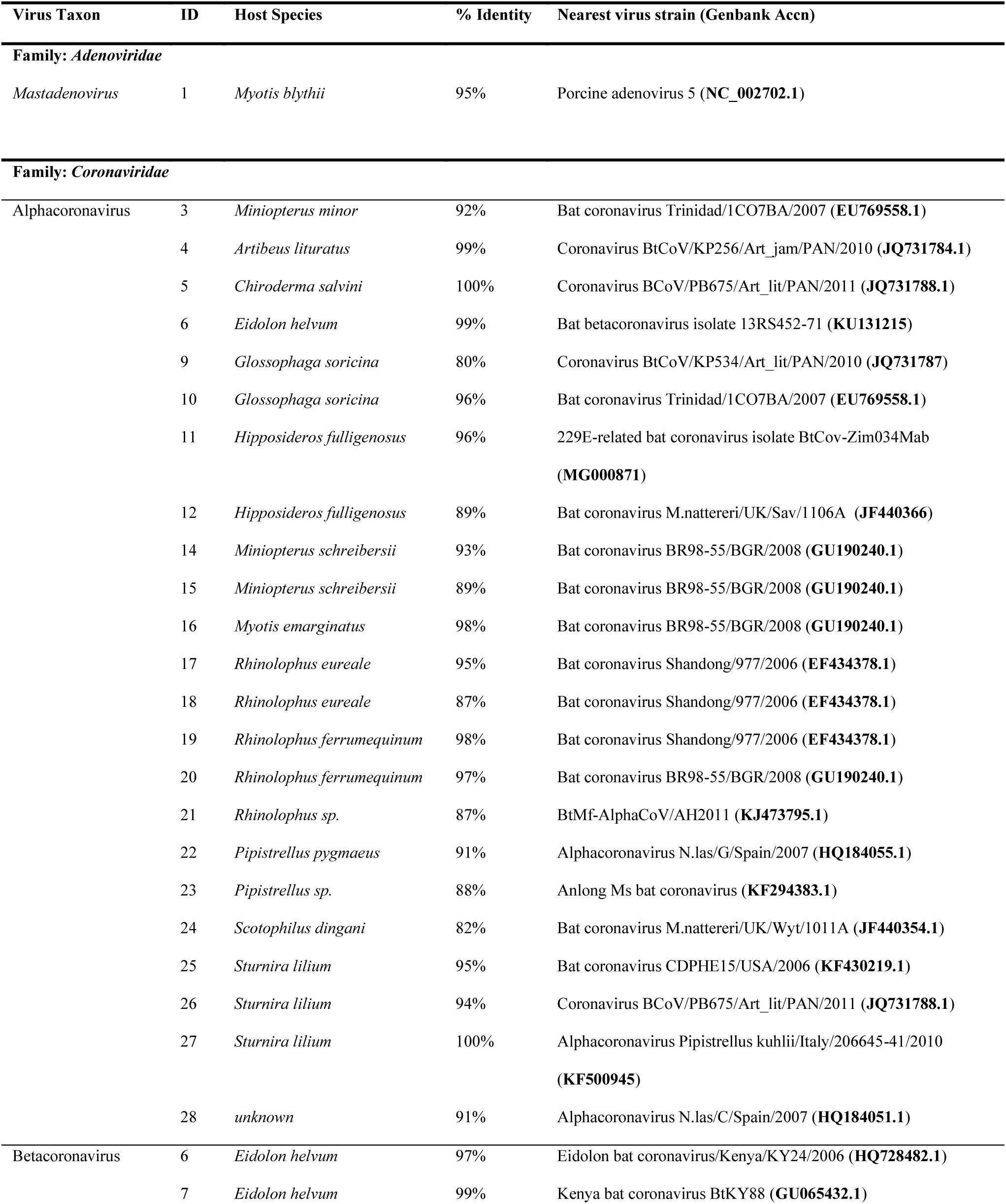
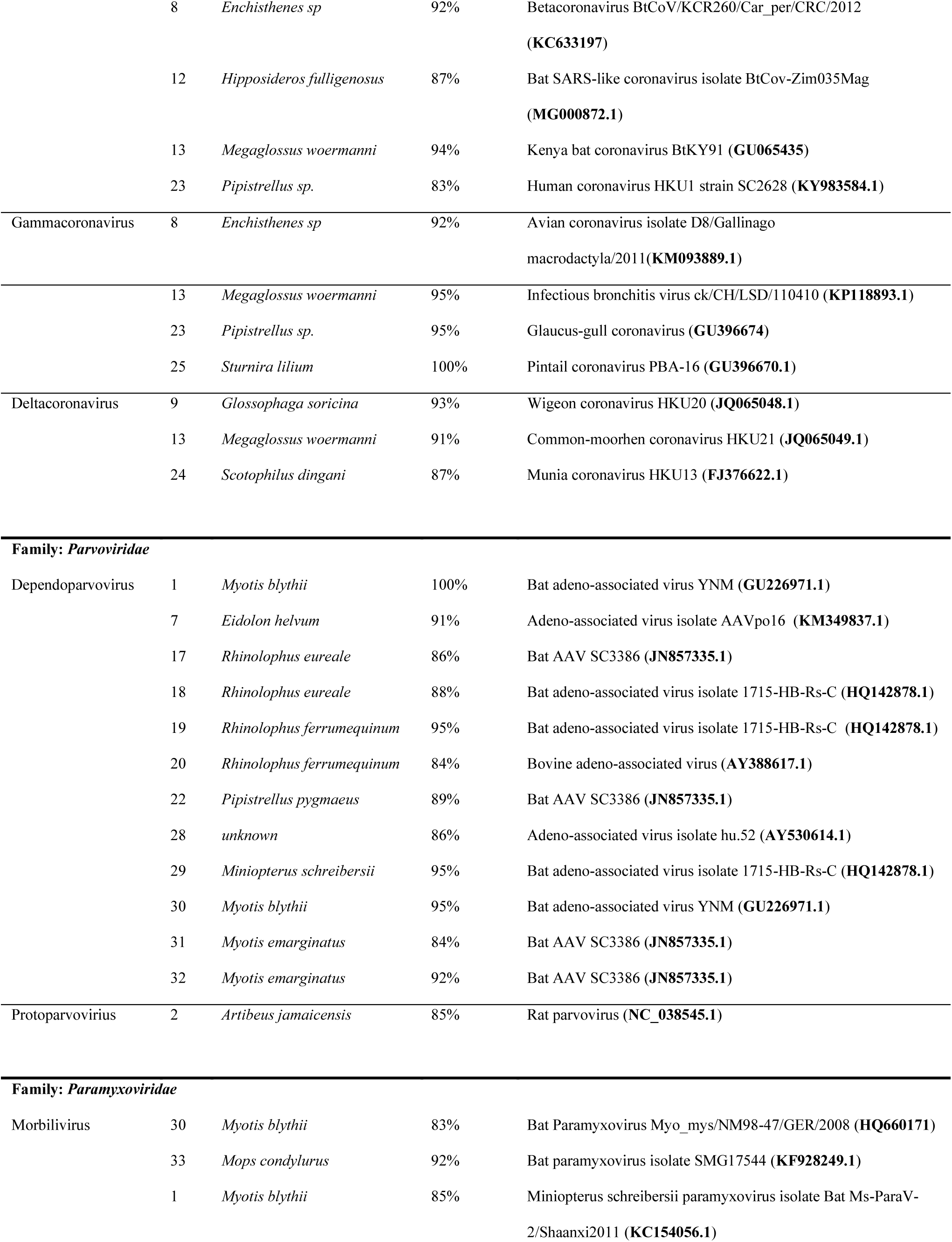
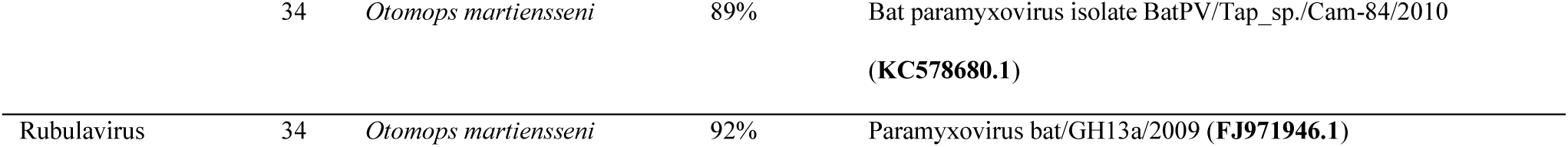
Viruses identified from bats. ID = individual bat identification number. % identity = BLAST nucleotide identity of amplicon to nearest neighbor

| Virus Taxon | ID | Host Species | % Identity | Nearest virus strain (Genbank Accn) |
| --- | --- | --- | --- | --- |
| **Family: Adenoviridae** |  |  |  |  |
| *Mastadenovirus* | 1 | *Myotis blythii* | 95% | Porcine adenovirus 5 (NC_002702.1) |
| **Family: Coronaviridae** |  |  |  |  |
| Alphacoronavirus | 3 | *Miniopterus minor* | 92% | Bat coronavirus Trinidad/1CO7BA/2007 (EU769558.1) |
|  | 4 | *Artibeus lituratus* | 99% | Coronavirus BtCoV/KP256/Art_jam/PAN/2010 (JQ731784.1) |
|  | 5 | *Chiroderma salvini* | 100% | Coronavirus BCoV/PB675/Art_lit/PAN/2011 (JQ731788.1) |
|  | 6 | *Eidolon helvum* | 99% | Bat betacoronavirus isolate 13RS452-71 (KU131215) |
|  | 9 | *Glossophaga soricina* | 80% | Coronavirus BtCoV/KP534/Art_lit/PAN/2010 (JQ731787) |
|  | 10 | *Glossophaga soricina* | 96% | Bat coronavirus Trinidad/1CO7BA/2007 (EU769558.1) |
|  | 11 | *Hipposideros fulligenosus* | 96% | 229E-related bat coronavirus isolate BtCov-Zim034Mab (MG000871) |
|  | 12 | *Hipposideros fulligenosus* | 89% | Bat coronavirus M.nattereri/UK/Sav/1106A (JF440366) |
|  | 14 | *Miniopterus schreibersii* | 93% | Bat coronavirus BR98-55/BGR/2008 (GU190240.1) |
|  | 15 | *Miniopterus schreibersii* | 89% | Bat coronavirus BR98-55/BGR/2008 (GU190240.1) |
|  | 16 | *Myotis emarginatus* | 98% | Bat coronavirus BR98-55/BGR/2008 (GU190240.1) |
|  | 17 | *Rhinolophus eurae* | 95% | Bat coronavirus Shandong/977/2006 (EF434378.1) |
|  | 18 | *Rhinolophus eurae* | 87% | Bat coronavirus Shandong/977/2006 (EF434378.1) |
|  | 19 | *Rhinolophus ferrumequinum* | 98% | Bat coronavirus Shandong/977/2006 (EF434378.1) |
|  | 20 | *Rhinolophus ferrumequinum* | 97% | Bat coronavirus BR98-55/BGR/2008 (GU190240.1) |
|  | 21 | *Rhinolophus sp.* | 87% | BtMf-AlphaCoV/AH2011 (KJ473795.1) |
|  | 22 | *Pipistrellus pygmaeus* | 91% | Alphacoronavirus N.las/G/Spain/2007 (HQ184055.1) |
|  | 23 | *Pipistrellus sp.* | 88% | Anlong Ms bat coronavirus (KF294383.1) |
|  | 24 | *Scotophilus dingani* | 82% | Bat coronavirus M.nattereri/UK/Wyt/1011A (JF440354.1) |
|  | 25 | *Sturnira lilium* | 95% | Bat coronavirus CDPHE15/USA/2006 (KF430219.1) |
|  | 26 | *Sturnira lilium* | 94% | Coronavirus BCoV/PB675/Art_lit/PAN/2011 (JQ731788.1) |
|  | 27 | *Sturnira lilium* | 100% | Alphacoronavirus Pipistrellus kuhlii/Italy/206645-41/2010 (KF500945) |
|  | 28 | *unknown* | 91% | Alphacoronavirus N.las/C/Spain/2007 (HQ184051.1) |
| Betacoronavirus | 6 | *Eidolon helvum* | 97% | Eidolon bat coronavirus/Kenya/KY24/2006 (HQ728482.1) |
|  | 7 | *Eidolon helvum* | 99% | Kenya bat coronavirus BtKY88 (GU065432.1) |
|  | 8 | *Enchisthenes sp* | 92% | Betacoronavirus BtCoV/KCR260/Car_per/CRC/2012<br>(**KC633197**) |
|  | 12 | *Hipposideros fulligenosus* | 87% | Bat SARS-like coronavirus isolate BtCov-Zim035Mag<br>(**MG000872.1**) |
|  | 13 | *Megaglossus woermanni* | 94% | Kenya bat coronavirus BtKY91 (**GU065435**) |
|  | 23 | *Pipistrellus sp.* | 83% | Human coronavirus HKU1 strain SC2628 (**KY983584.1**) |
| Gammacoronavirus | 8 | *Enchisthenes sp* | 92% | Avian coronavirus isolate D8/Gallinago<br>macroductyla/2011(**KM093889.1**) |
|  | 13 | *Megaglossus woermanni* | 95% | Infectious bronchitis virus ck/CH/LSD/110410 (**KP118893.1**) |
|  | 23 | *Pipistrellus sp.* | 95% | Glaucus-gull coronavirus (**GU396674**) |
|  | 25 | *Sturnira lilium* | 100% | Pintail coronavirus PBA-16 (**GU396670.1**) |
| Deltacoronavirus | 9 | *Glossophaga soricina* | 93% | Wigeon coronavirus HKU20 (**JQ065048.1**) |
|  | 13 | *Megaglossus woermanni* | 91% | Common-moorhen coronavirus HKU21 (**JQ065049.1**) |
|  | 24 | *Scotophilus dingani* | 87% | Munia coronavirus HKU13 (**FJ376622.1**) |
| **Family: Parvoviridae** |  |  |  |  |
| Dependoparvovirus | 1 | *Myotis blythii* | 100% | Bat adeno-associated virus YNM (**GU226971.1**) |
|  | 7 | *Eidolon helvum* | 91% | Adeno-associated virus isolate AAVpo16 (**KM349837.1**) |
|  | 17 | *Rhinolophus eurae* | 86% | Bat AAV SC3386 (**JN857335.1**) |
|  | 18 | *Rhinolophus eurae* | 88% | Bat adeno-associated virus isolate 1715-HB-Rs-C (**HQ142878.1**) |
|  | 19 | *Rhinolophus ferrumequinum* | 95% | Bat adeno-associated virus isolate 1715-HB-Rs-C (**HQ142878.1**) |
|  | 20 | *Rhinolophus ferrumequinum* | 84% | Bovine adeno-associated virus (**AY388617.1**) |
|  | 22 | *Pipistrellus pygmaeus* | 89% | Bat AAV SC3386 (**JN857335.1**) |
|  | 28 | *unknown* | 86% | Adeno-associated virus isolate hu.52 (**AY530614.1**) |
|  | 29 | *Miniopterus schreibersii* | 95% | Bat adeno-associated virus isolate 1715-HB-Rs-C (**HQ142878.1**) |
|  | 30 | *Myotis blythii* | 95% | Bat adeno-associated virus YNM (**GU226971.1**) |
|  | 31 | *Myotis emarginatus* | 84% | Bat AAV SC3386 (**JN857335.1**) |
|  | 32 | *Myotis emarginatus* | 92% | Bat AAV SC3386 (**JN857335.1**) |
| Protoparvovirus | 2 | *Artibeus jamaicensis* | 85% | Rat parvovirus (**NC_038545.1**) |
| **Family: Paramyxoviridae** |  |  |  |  |
| Morbilivirus | 30 | *Myotis blythii* | 83% | Bat Paramyxovirus Myo_mys/NM98-47/GER/2008 (**HQ660171**) |
|  | 33 | *Mops condylurus* | 92% | Bat paramyxovirus isolate SMG17544 (**KF928249.1**) |
|  | 1 | *Myotis blythii* | 85% | Miniopterus schreibersii paramyxovirus isolate Bat Ms-ParaV-2/Shaanxi2011 (**KC154056.1**) |
|  | 34 | *Otomops martiensseni* | 89% | Bat paramyxovirus isolate BatPV/Tap_sp./Cam-84/2010<br>(**KC578680.1**) |
| Rubulavirus | 34 | *Otomops martiensseni* | 92% | Paramyxovirus bat/GH13a/2009 (**FJ971946.1**) |

## 4. Discussion

A sensitive and high throughput identification system for a broad range of viruses is needed in cases of unknown viral disease outbreaks in humans or animals. A major challenge is that viruses lack a shared universal phylogenetic marker such as 16S rRNA for bacteria or 18S rRNA for eukaryotes. AA-PCR-Seq provides a solution to identify a wide range of known and potentially novel viruses from large, complex sample sets. Using a respiratory virus panel as a proof of concept, we evaluated AA-PCR-Seq as a comprehensive virus detection and discovery platform, comparing its relative sensitivity to shotgun NGS as well as to qPCR and conventional pan-viral group PCR. Finally, we used AA-PCR-Seq to test a large set of human clinical samples as well as samples from bats, and we were able to detect a wide array of viruses with varying degrees of homology to known species.

Shotgun NGS, PCR/qPCR, and AA-PCR-Seq are all useful and valid tools in virus detection and discovery, each with certain strengths and weaknesses. Next-generation sequencing allows for detection of a very broad array of pathogens, but it lacks the sensitivity, scalability, and low cost of PCR. On the other hand, though relatively inexpensive, sensitive, and straightforward, PCR strategies can be too narrowly targeted to identify a large panel of viruses or novel viruses. Unlike qPCR, consensus degenerate pan-viral group PCR provides broad specificity to amplify a broad range of known viruses as well as distantly related virus species. AA-PCR-Seq builds upon the strength of both NGS and pan-viral group PCR, using microfluidics to run dozens of these assays in parallel on dozens of samples. NGS allows analysis of these thousands of amplicons in a fraction of the time, effort, and cost it otherwise would take to purify and sequence individual amplicons using gels and a capillary sequencer.

Pan-viral group PCR assays are, by design, much more broadly-reactive than qPCR. Because of that design choice, and the small sample volume afforded by the Access Array, some AA-PCR-Seq assays were initially less sensitive compared to full-volume, traditional PCR. However, we were able to compensate for much of the loss in sensitivity by incorporating a pre-amplification step, which increases the concentration of total nucleic acid entering each AA-PCR-Seq reaction. Because of the ability to test up to 48 samples concurrently using up to 48 viral assays we believe the sensitivity level of AA-PCR-Seq is acceptable and within the Ct range of most clinical samples. In addition, the assays included in AA-PCR-Seq are not limited to detecting known viruses. We demonstrate this case with zoonotic virus testing where AA-PCR-Seq was able to detect 55 diverse and potentially novel viruses in the bat panel without prior knowledge of their sequences. There are no existing qPCR primers available for these viruses and many of them were not detectable by direct shotgun sequencing..

AA-PCR-Seq does not require multiplexing optimization, since it uses singelplex PCR assays in individual reaction chambers. Further, the Access Array is modular so that it is simple to add or replace assays. This is especially useful because pan-viral group PCR primers are updated and improved as new viral sequences become available. In addition to the respiratory panel described in this manuscript, we have also added a full complement of pan-viral group PCR assays for two main syndrome diseases including gastroenteritis and encephalitis. Additional viruses were identified from the bat sample set, including two astroviruses, four orthoreoviruses, and three rotaviruses. Work is still being done to fully validate these expanded panels.

Finally, AA-PCR-Seq increases sample throughput while reducing reagent and sample usage. Excluding capital costs, when run at capacity in screening large sample sets, the reagent cost of each AA-PCR-Seq sample/assay pair is reduced 10-50-fold compared to conventional PCR and Sanger sequencing. Furthermore, the microfluidic system requires a small volume of sample (<10 µL) to test for the entire panel of viruses, whereas conventional pan-viral group PCR would require nearly 200 µL of sample. This is critical during unknown case investigations when sample volume is limited for screening by a large panel of assays for differential testing.

We see AA-PCR-Seq as a strategy that complements PCR and shotgun NGS strategies. With exponentially more sequencing effort (and the associated cost, time, and data analysis and storage concerns), shotgun NGS should be able to identify all the viruses in this study and more. However, labs that can afford routine NGS frequently rely on the MiSeq, which is a lower output instrument. AA-PCR-Seq enables a very efficient use of these reads and facilitates use of the MiSeq as a high throughput broad virus detection and discovery platform. Similarly, AA-PCR-Seq is not meant to supplant qPCR or function as a clinical diagnostic. Its strength is in screening large number of samples from unknown outbreaks and surveillance studies, providing more broad detection capabilities than is afforded by qPCR/PCR without the cost of massive high-throughput shotgun sequencing.

## Data statement

The sequence data generated in this study is deposited in the NCBI SRA under BioProjects **PRJNA573045** and **PRJNA574475**.

## Acknowledgments

We wish to acknowledge the Centers for Disease Control and Prevention Office of Public Health Preparedness and Response as well as the Office of Advanced Molecular Detection for help in funding this project. We also wish to thank Dr. Anne Whitney and Dr. Michael Farrell for helpful comments and discussion early in this project.

